# TAGINE: Fast Taxonomy-based Feature Engineering for Microbiome Analysis

**DOI:** 10.1101/2025.11.25.689983

**Authors:** Shiri Baum, Ido Meshulam, Yadid M Algavi, Omri Peleg, Elhanan Borenstein

## Abstract

**Summary:** TAGINE is a feature engineering algorithm that leverages the microbial taxonomic tree to optimize feature sets in microbiome data for predictive modeling. The algorithm starts with features at high taxonomic levels and iteratively splits them into lower-level clades in cases where it improves predictive accuracy, ultimately producing a feature set spanning multiple taxonomic levels. This approach aims to markedly reduce the number of features while preserving biological relevance and interpretability. We compare TAGINE’s performances to other standard and taxonomy-based feature engineering methods on several different datasets, and show that TAGINE yields more compact feature sets and is orders of magnitude faster than other methods, while maintaining predictive accuracy.

**Availability and Implementation:** TAGINE is freely available under the MIT license with source code available at https://github.com/borenstein-lab/tagine_fe.

## Introduction

Recent advances in sequencing technologies have facilitated the high-throughput and high-resolution acquisition of biomedical data (Soon *et al*., 2013). These developments offer tremendous promise for improving disease diagnosis and treatment (Schupack *et al*., 2022), but also introduce considerable analytical, computational, and conceptual challenges. Perhaps the most daunting challenge when analyzing such high-dimensional data, where the number of features often vastly exceeds the number of samples (Bellman and Kalaba, 1959), is the risk of identifying spurious associations or overfitting predictive model to the training data. Traditional feature selection and engineering methods aim to mitigate this issue by pinpointing highly informative features or by combining multiple features into composite ones using a variety of statistical or computational approaches (Kuzudisli *et al*., 2023). While these methods provide an effective strategy for many analytical tasks (including the construction of machine-learning predictive models), they often overlook existing biological knowledge, thus limiting model interpretability, failing to utilize available information, and reducing their utility for hypothesis generation.

One notable domain that is characterized by both high-dimensional data and extensive prior biological knowledge is microbiome research. Metagenomics-based profiling of various microbiomes often generates datasets comprising thousands to tens of thousands of species-level taxonomic features (with most studies including only a few dozen to several hundred samples). On the other hand, taxonomic classification systems and phylogenetic trees that link the various taxa offer a natural way to group different taxonomic features. Moreover, since phylogenetic proximity often reflects functional or ecological similarity, such phylogeny- or taxonomy-based grouping is likely analytically and biologically relevant (Goberna and Verdu, 2018).

With that in mind, many microbiome studies indeed aggregate features at some coarse taxonomic level (e.g., genus or phylum) prior to downstream analyses in the hope of reducing the number of taxonomic features and improving statistical power (Kleine Bardenhorst *et al*., 2021). However, such fixed-level aggregation may obscure clade-specific differences in granularity and information content, potentially overlooking highly informative features that reside at different taxonomic levels. To address this limitation, a small number of recently introduced methods have introduced tree-guided adaptive grouping of features, aiming to combine informative fine-grained taxonomic features in some clades with coarser-grained features in others. Trac (Bien *et al*., 2021), for example, focuses on regression tasks, using convex optimization to jointly learn predictive groupings and model coefficients. Other methods focus, as do we, on classification tasks: HFE (*Hierarchical Feature Engineering*) (Oudah and Henschel, 2018) iteratively identifies informative features along paths in the taxonomic tree, ultimately providing a set of features at various taxonomic levels along these paths, while TaxaHFE (Oliver *et al*., 2023), utilizes an iterative machine learning approach to collapse information-poor features into higher levels. These classification-focused methods were shown to exhibit impressive predictive performances and feature reduction, while preserving relevant community structure. Importantly, however, these methods are computationally heavy and suffer from relatively slow runtimes, which markedly limit their applicability in large-scale settings. Furthermore, some of these methods can also include multiple features on the same tree path in the final feature set, limiting interpretability.

Motivated by these observations, we introduce TAGINE (Taxonomy-Aware Grouping for INformation Enhancement), a new and fast feature engineering algorithm that leverages the taxonomical tree hierarchy to guide dimensionality reduction for classification tasks. TAGINE begins with coarse-grained taxonomic features and iteratively splits them into finer-grained levels, prioritizing splits that improve predictive accuracy. The resulting grouped features can be then used as inputs for downstream analysis or predictive modeling. To benchmark TAGINE’s effectiveness, we compared its performance to recursive feature elimination (RFE) (Guyon *et al*., 2002), a commonly used, domain agnostic feature selection algorithm, as well as two of the previously noted taxonomy-based hierarchical feature engineering algorithms. We evaluated the predictive accuracy obtained when using the resulting feature sets, the sizes of these sets, and the runtimes of each method on several independent datasets, showing that TAGINE significantly reduces the number of features and improves runtime while maintaining predictive accuracy.

### The TAGINE Algorithm

The TAGINE algorithm takes as input a dataset of taxonomic abundance values, along with a corresponding taxonomic tree structure *T*, such that each feature in the dataset corresponds to a leaf in *T*. Internal nodes of *T* represent higher-level taxonomic feature, with their corresponding abundance values computed as the sum of their children’s abundances. TAGINE further defines a queue, *Q*, which contains nodes that are candidates for expansion and that will be evaluated iteratively by the algorithm.

TAGINE begins with the tree collapsed up to a predefined taxonomic level (by default, one level below the root), leaving a small number of coarse-grained features, which are subsequently added to *Q*. TAGINE then iteratively pops a collapsed node from *Q*, and compares two models to determine whether this node should be expanded: one that treats the node as a single feature to predict the label, and the other that uses the node’s children as separate features. Specifically, a logistic regression is fitted to each of the two models, and the quality of each model is determined using the Akaike Information Criterion (AIC; Akaike, 1974). If the AIC suggests that the single-parameter model is a better fit, this means that expanding the node does not improve prediction, and the node remains collapsed as a single feature in the final feature set (which also means its descendants will not be examined by the algorithm). Otherwise, the node is expanded, and any child that represents a collapsed node is added to *Q* for further evaluation. In the implementation analyzed below, to further reduce the number of features, features whose coefficient in the logistic regression was not significant were pruned from *T* (and not added to *Q*).

The algorithm ends when the queue is empty, and the features that remain in *T* as leaves (which may include features at various taxonomical levels, including collapsed nodes) are used as the final feature set.

## Evaluation and Results

To evaluate the performance of TAGINE, we applied it to four shotgun metagenomic datasets from the Microbiome-Metabolome curated resource (Muller *et al*., 2022). The datasets included studies of inflammatory bowel disease (IBD; Franzosa et al., 2019), colorectal cancer (CRC; Yachida et al., 2019), end-stage renal disease (ESRD; Wang et al., 2020), and gastric cancer (GC; Erawijantari et al., 2020). For each dataset, we randomly partitioned the data into 50 training and test sets using an 85%-15% split. We then removed very rare species from each dataset (mean abundance below 10^−6^ or mean prevalence below 10^−2^ in the training set), leaving a mean of 6943.5, 7081.4, 6824.0, and 7121.6 species-level features in the CRC, ESRD, GC, and IBD datasets, respectively. After filtration, the data was normalized to represent relative abundances. Finally, to address zero values, a small constant equal to half the minimum value in each sample was added to all features in that sample, followed by renormalization.

Given these data, we applied TAGINE to each training set (recording the runtime and the resulting number of features), used the selected features to train a random forest model on a classification task, applied this classifier to the test samples, and computed the area under the Receiver Operating Characteristic curve (AUROC; Bradley, 1997). As a classification task, we utilized the ESRD and GC datasets in their original case-control format (i.e., classifying cases vs. controls). For the CRC dataset, we classified advanced-stage CRC versus all other categories, and for the IBD dataset, we classified UC/CD versus controls.

We compared TAGINE to four other feature selection/engineering methods: (i) No feature selection, (ii) Scikit-learn’s (Pedregosa *et al*., 2011) recursive feature elimination algorithm, applied to the data after all possible taxonomical levels are included and configured to select the same number of features as TAGINE, (iii) HFE (Oudah and Henschel, 2018), and (iv) TaxaHFE (Oliver *et al*., 2023). TaxaHFE was specifically used with the super filter feature, as it provided fewer features and better predictive performance.

We first examined the predictive accuracy obtained with the set of features selected by each approach (Figure 1A). We found that all methods had comparable performance on most datasets. In fact, in only one of the four datasets (ESRD), was the AUROC of the best performing method (RFE) significantly higher than that of TAGINE (*p* = 4 · 10^−3^, one-sided *t* test with FDR corrected *p* < 0.05). In the CRC dataset, TAGINE achieved significantly higher AUROC scores than two of the other methods (*p* = 0.03 and *p* = 1.6 · 10^−5^ for no feature selection and RFE, respectively).

**Figure 1.**
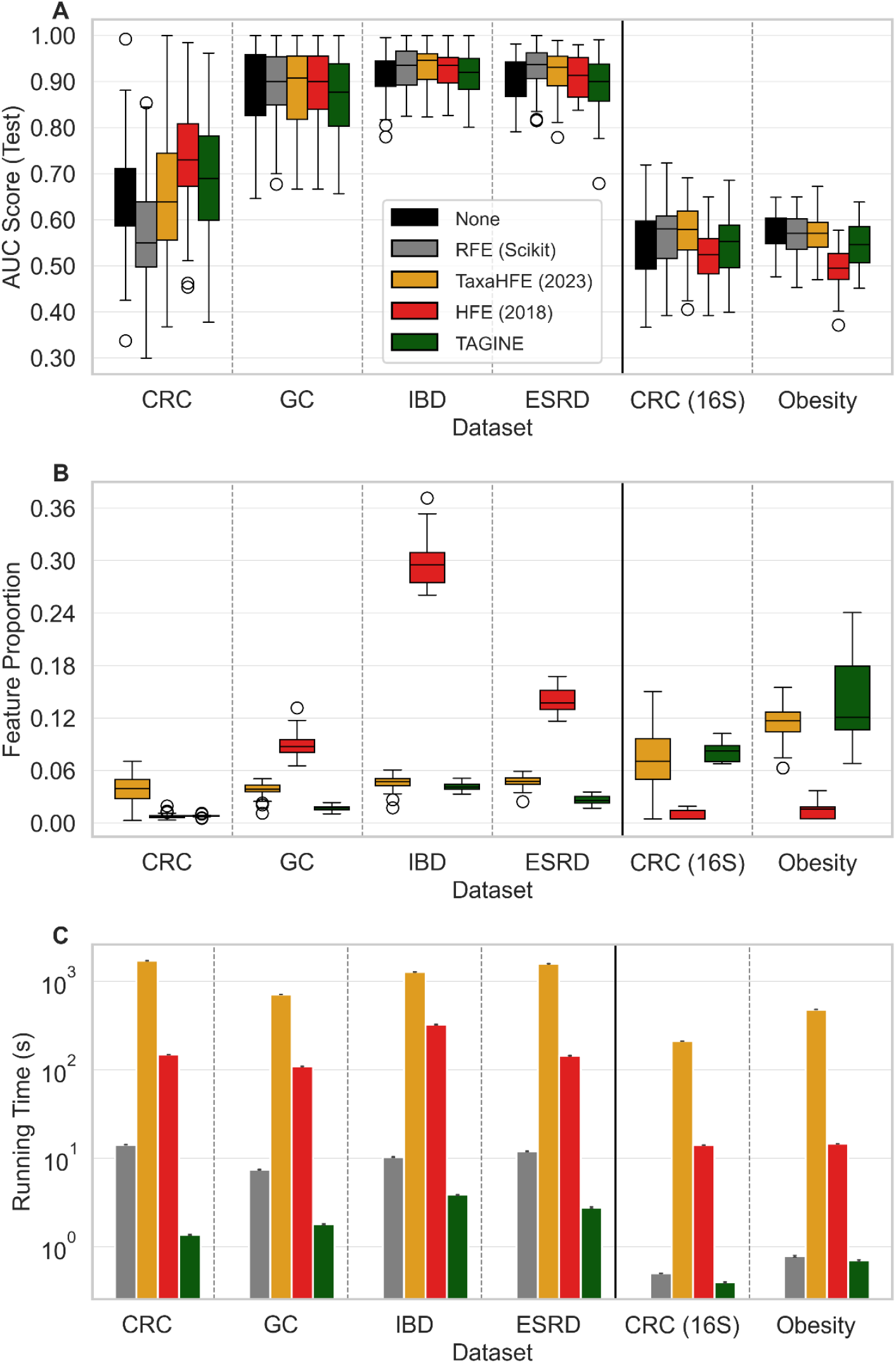
(A) Comparison of AUC scores by dataset and selection method. (B) Comparison of the proportion of features selected out of the original set of features by dataset and selection method. (C) Comparison of running time by dataset and selection method.

We then turned to evaluating the number of features selected by each method (Figure 1B). For this analysis, we naturally only compared TAGINE to HFE and TaxaHFE, since RFE was preconfigured to select the same number of features as TAGINE. We found that in terms of number of features, TAGINE significantly outperformed both tested methods in all datasets but one (in the CRC dataset, HFE outperformed TAGINE, though the difference was not statistically significant with *p* = 1 after FDR correction). It is also worth noting the incredibly effective reduction in the number of features selected by TAGINE, with the size of the final feature set being only 0.8%, 2.7%, 1.7%, and 4.1% of the total number of features in the CRC, ESRD, GC, and IBD datasets, respectively. This is particularly impressive given our finding above that the predictive accuracy achieved using these feature sets was comparable, and sometimes exceeded, the accuracy achieved using the full set of features.

Perhaps the biggest advantage of TAGINE is its simplicity and, as a result, its short running time. Indeed, comparing TAGINE to all other methods (except for the no-feature-selection baseline), we found that it was significantly and substantially faster than all other methods in all four datasets (Figure 1C). For example, in the smallest dataset (GC, with a mean of 6824.0 initial features), TAGINE ran, on average, 4.2 times faster than RFE, 61.1 times faster than HFE, and 395.4 times faster than TaxaHFE (average runtimes 1.8s for TAGINE, 7.5s for RFE, 109.9s for HFE, and 711.7s for TaxaHFE, measured on a 13^th^ Gen Intel® Core™ i7 processor). This advantage persisted in largest datasets, with TAGINE running 2.7, 85.8, and. 337.3 times faster than RFE, HFE, and

TaxaHFE, respectively, on the largest IBD dataset (average runtimes 3.9s for TAGINE, 10.3s for RFE, 326.0s for HFE, and 1282.8s for TaxaHFE). This reduction in runtime is of great practical relevance, as it makes TAGINE a viable option for large-scale datasets or for tasks requiring multiple iterations, where other methods may prove too computationally demanding to apply.

Finally, to evaluate TAGINE’s performances on substantially smaller datasets, we applied it to two 16S rRNA cohorts from the Microbiome HD collection (Duvallet *et al*., 2017), including a study of CRC (Baxter *et al*., 2016) and obesity (Goodrich *et al*., 2014). The data was processed as described above, and the same feature selection and engineering methods were evaluated. Notably, the two datasets are characterized by a much smaller initial number of features (a mean of 205.0 for CRC, and 189.1 for obesity). On these data, TAGINE still significantly outperformed all other methods in terms of runtime, but was often outperformed in terms of number of features and predictive performance by some other methods, suggesting that TAGINE’s primary advantage is more pronounced on larger datasets.

## Conclusion

We present TAGINE, a fast, taxonomy-based feature engineering algorithm that substantially outperforms existing methods in terms of speed and the number of features selected, while maintaining similar predictive performance. TAGINE excels specifically in large datasets with a large number of features, which, naturally, are precisely the cases where feature selection and engineering methods are most needed.

TAGINE’s short runtime makes it a flexible and cost-effective processing step, which can be easily included in various data processing, analysis, or machine learning pipelines. Furthermore, its fast runtime on large datasets mean that it does not necessitate aggressive filtering of rare taxa, which may be required prior to the application of other methods. This may prove crucial in datasets where this kind of prevalence or abundance filtering obscures the signal, ultimately degrading predictive performance. By using TAGINE, large initial feature sets can be retained without incurring significant runtime or performance costs. We further note that TAGINE can be used on any hierarchal data, expanding its applicability beyond microbiome studies.

Importantly, each feature selected by TAGINE corresponds to a single clade in the taxonomic tree (without nesting across different levels), offering a highly interpretable feature set, especially compared to more domain-agnostic methods. In addition to selecting a set of features for downstream predictive modelling, this property of TAGINE also means that it yields a pruned taxonomic tree from which these features are derived. We propose that this pruned tree may, in and of itself, serve as a useful interpretative tool for exploring disease-specific microbial patterns. For instance, comparing the topologies of pruned trees across related vs unrelated diseases could reveal shared vs. divergent microbial signatures. Moreover, examining which taxa are consistently retained or excluded across disease contexts may help identify species or clades that exhibit disease-specific variability.

## Acknowledgements

A preliminary version of this method was developed as part of a workshop in microbiome analysis, mentored by Yadid M Algavi and Omri Peleg. We would like to thank all the participants of this workshop, as well as members of the Borenstein lab, for helpful feedback and insightful discussions.

## Funding

This work was supported, in part, by Israel Science Foundation (ISF) grant 2435/19, by Len Blavatnik and the Blavatnik Family foundation, and by the Koret-UC Berkeley-Tel Aviv University Initiative in Computational Biology and Bioinformatics to E.B. SB and OP are supported, in part, by a fellowship from the Edmond J. Safra Center for Bioinformatics at Tel-Aviv University. The funders had no role in study design, data collection and analysis, decision to publish, or preparation of the manuscript.

## Notes

### Competing Interest Statement

The authors have declared no competing interest.

